# Characterisation of the *ERF102* to *ERF105* genes of *Arabidopsis thaliana* and their role in the response to cold stress

**DOI:** 10.1101/848705

**Authors:** Sylvia Illgen, Stefanie Zintl, Ellen Zuther, Dirk K. Hincha, Thomas Schmülling

**Affiliations:** Institute of Biology/Applied Genetics, Dahlem Centre of Plant Sciences (DCPS), Freie Universität Berlin, D-14195 Berlin, Germany; Max-Planck-Institute of Molecular Plant Physiology, D-14476 Potsdam, Germany

**Keywords:** *Arabidopsis thaliana*, cold acclimation, *ETHYLENE RESPONSE FACTOR* genes, freezing tolerance, root architecture, transcription factor

## Abstract

The *ETHYLENE RESPONSE FACTOR* (*ERF*) genes of *Arabidopsis thaliana* form a large family encoding plant-specific transcription factors. Here, we characterise the four phylogenetically closely related *ERF102/ERF5*, *ERF103/ERF6*, *ERF104* and *ERF105* genes. Expression analyses revealed that these four genes are similarly regulated by different hormones and abiotic stresses. Analyses of tissue-specific expression using *promoter:GUS* reporter lines revealed their predominant expression in root tissues including the root meristem (*ERF103*), the quiescent center (*ERF104*) and the root vasculature (all). All GFP-ERF fusion proteins were nuclear-localised. The analysis of insertional mutants, amiRNA lines and *35S:ERF* overexpressing transgenic lines indicated that *ERF102* to *ERF105* have only a limited impact on regulating shoot and root growth. Previous work had shown a role for ERF105 in the cold stress response. Here, measurement of electrolyte leakage to determine leaf freezing tolerance and expression analyses of cold-responsive genes revealed that the combined activity of ERF102 and ERF103 is also required for a full cold acclimation response likely involving the CBF regulon. Together, these results suggest a common function of these *ERF* genes in regulating root architecture and the response to cold stress.

## INTRODUCTION

The *ERF* genes encode plant-specific transcription factors forming a large gene family with 122 members in *Arabidopsis thaliana* (Nakano, Suzuki, Fujimura & Shinshi, 2006). The ERF transcription factors are members of the APETALA2/ETHYLENE RESPONSE FACTOR (AP2/ERF) superfamily, which also contains the AP2 and RAV families and which is defined by the AP2/ERF DNA-binding domain (Riechmann *et al*., 2000). This domain is about 60 amino acids long and forms an interface of three antiparallel β-strands and one α-helix (Ohme-Takagi & Shinshi, 1995). The β-strands bind to an 11 bp consensus sequence (5’-TAAGAGCCGCC-3’), the GCC-Box, in the major groove of the DNA double helix (Hao, Ohme-Takagi & Sarai, 1998). ERF transcription factors are involved in the regulation of numerous developmental processes (Riechmann & Meyerowitz, 1998) and they are important for the response to various biotic and abiotic stresses including cold (Agarwal, Agarwal, Reddy & Sopory, 2006b; Kizis, Lumbreras & Pages, 2001; Srivastava & Kumar 2019; Xie, Nolan, Jiang & Yin, 2019).

Previously, we identified four phylogenetically closely related *ERF* genes with similar transcriptional responses to cytokinin (Brenner, Romanov, Köllmer, Bürkle & Schmülling, 2005). These genes, *ERF102* (AT5G47230; known as *ERF5*), *ERF103* (AT4G17490; identical to *ERF6*), *ERF104* (AT5G61600) and *ERF105* (AT5G51190) are members of group IXb of the ERF family (Nakano *et al*., 2006). Expression of *ERF102* to *ERF105* is regulated by cold and different cold stress-related hormones, and it was demonstrated that *ERF105* has a function in the freezing tolerance and cold acclimation of *Arabidopsis* (Bolt, Zuther, Zintl, Hincha & Schmülling, 2017). All four *ERF* genes are also involved in the response to other stresses*. ERF102* and *ERF103* regulate leaf growth inhibition upon mild osmotic stress (Dubois *et al*., 2013, 2015) and *ERF103* additionally regulates oxidative stress responses (Sewelam *et al*., 2013). *ERF103*, *ERF104* and *ERF105* are involved in the fast retrograde signalling response and the acclimation response to high light (Moore, Vogel & Dietz, 2014; Vogel *et al*., 2014). Further studies have shown that *ERF102* to *ERF105* play a role in plant immunity (Bethke *et al*., 2009; Cao *et al*., 2019; Mase *et al*., 2013; Meng *et al*., 2013; Moffat *et al*., 2012; Son *et al*., 2012). Thus, ERF102 to ERF105 match the profile of other ERF transcription factors designated as a regulatory hub integrating hormone signalling in the plant response to abiotic stresses (Müller & Munné-Bosch, 2015).

The close phylogenetic relationship among the four *ERF* genes and the similarity of their transcriptional responses to different cues suggested that they share some common functions in response to cold. Cold stress adversely affects plant growth and development and several pathways to respond to cold stress have been described. Plants from temperate and boreal climates have evolved mechanisms to acquire freezing tolerance through cold acclimation, a process in which upon exposure to low non-freezing temperatures the ability to survive freezing temperatures increases (Xin & Browse, 2000). A central cold signalling pathway is the CBF (*C-REPEAT-BINDING FACTOR/DEHYDRATION-RESPONSE ELEMENT-BINDING PROTEIN*) regulon. The *CBF1* (*DREB1b*), *CBF2* (*DREB1c*) and *CBF3* (*DREB1a*) genes are the central regulatory elements of this regulon (Chinnusamy, Zhu & Zhu, 2007; Liu *et al*., 1998). The INDUCER OF C-REPEAT-BINDING FACTOR EXPRESSION 1 (ICE1), a MYC-type bHLH (basic helix-loop-helix) transcription factor, is post-translationally activated in response to cold (Chinnusamy *et al*., 2003; Ding *et al*., 2015; Li *et al*., 2017; Miura *et al*., 2007). ICE1 in turn activates the transcription of the *CBF3* gene (Chinnusamy *et al*., 2003). Besides ICE1, expression of the cold-regulated *CBF* genes is positively controlled by several other transcription factors including ICE2 and CALMODULIN-BINDING TRANSCRIPTION ACTIVATOR 3 (CAMTA3) (Doherty, Van Buskirk, Myers & Thomashow, 2009; Fursova, Pogorelko & Tarasov, 2009). Negative regulators of the CBF regulon are, for instance, the C2H2 zinc finger transcription factor ZAT12 (Vogel, Zarka, Van Buskirk, Fowler & Thomashow, 2005) and MYB15 (Agarwal *et al*., 2006a). MYB15 is in turn negatively regulated by ICE1 (Agarwal *et al*., 2006a) and phosphorylation of MYB15 by MPK6 reduces its affinity to bind to the *CBF3* promoter (Kim *et al*., 2017). The CBF proteins regulate the expression of the *COLD-REGULATED* (*COR*) genes and physiological responses (e.g. accumulation of cryoprotective compounds, modification of cellular structures) that together confer cold acclimation (Thomashow, 1999; Yamaguchi-Shinozaki & Shinozaki, 2006). Transcriptomic analyses of the CBF regulon has revealed that only part (∼11%) of the cold-responsive genes is under control of the CBF regulon (Park *et al*., 2015), which was confirmed by gene expression analysis in *cfb* triple mutants (Jia *et al*., 2016; Zhao, Zhang, Xie, Si, Li & Zhu, 2016). It was concluded that only about one-third of the increase in freezing tolerance that occurs in response to low temperature is dependent on the CBF regulon (Park *et al*., 2015). Together, this suggests that an extensive regulatory network involving numerous transcription factors in addition to the best known CBF core regulators governs the response to cold.

We previously identified the *ERF105* gene of *Arabidopsis* as an important factor for *Arabidopsis* freezing tolerance and cold acclimation (Bolt *et al*., 2017). The strongly reduced expression of cold-responsive genes in *ERF105* mutants upon cold acclimation suggests that its action is linked to the CBF regulon. Also the expression of three closely related transcription factor genes, *ERF102*, *ERF103* and *ERF104*, is induced by cold (Bolt *et al*., 2017; Lee, Henderson & Zhua, 2005; Park *et al*., 2015; Vogel *et al*., 2005). It is therefore possible that these transcription factors have a function in the response to cold stress. Here, we have extended our analysis of the *ERF105* gene family. We provide additional transcript data supporting a similar response profile of the *ERF105* family members and show the tissue-specific expressions of *pERF102:GUS* to *pERF104:GUS* as well as the subcellular localisations of GFP-ERF102 to GFP-ERF104 fusion proteins. Single and combined loss-of-function mutants and lines overexpressing single *ERF* genes were analysed for their growth characteristics and cold stress response and reveal partial functional redundancy of the members of this transcription factor subfamily.

## MATERIAL AND METHODS

### Plant material

*Arabidopsis thaliana* accession Col-0 was used as wild type. The *erf105* mutant, *ERF105* overexpressing lines, *pERF105:GUS* lines, complementation lines of *erf105,* as well as 35S:ami104 and 35S:ami104/105 lines have been described previously (Bolt *et al*., 2017). The T-DNA insertion line *erf102* (SAIL_46_C02) was obtained from the Nottingham Arabidopsis Stock Centre (NASC). After selection of homozygous plants, the location of the T-DNA insertion was verified by sequencing and plants were backcrossed twice with Col-0 to eliminate possible multiple insertions and other background mutations. Complementation of the *erf102* phenotype was tested by introgressing ERF102ox-1 and ERF102ox-2 into the *erf102* background. To generate lines overexpressing *ERF102* to *ERF104*, the genomic coding sequences of *ERF102* to *ERF104* were amplified by PCR, cloned into pDONR221 (Invitrogen, Carlsbad, USA) by using the Gateway cloning system and transferred subsequently into vector pK7WGF2 (Karimi, Depicker & Hilson, 2007b). To generate *pERF102:GUS* to *pERF104:GUS* reporter genes, the promoter regions of the *ERF* genes (∼2 kb upstream of the start codon) were amplified by PCR and cloned into pDONR P4-P1R (Invitrogen). To generate the binary destination vectors, the pDONR P4-P1R constructs with the *ERF* promoters and the Gateway entry clone pEN-L1-SI-L2 (Karimi, Bleys, Vanderhaeghen & Hilson, 2007a) harboring the *GUS* reporter gene were then combined into the destination vector pK7m24GW,3 using MultiSite Gateway (Karimi, De Meyer & Hilson, 2005). Artificial microRNA (amiRNA) was used to generate lines with a reduced *ERF103* expression (Schwab, Ossowski, Riester, Warthmann & Weigel, 2006). amiRNAs directed against *ERF104* and *ERF105* were described (Bolt *et al*., 2017). The amiRNA sequence targeting *ERF103* was 5′-TAACGTCGTAACTTTCCCCCG-3′. The sequence was selected and the expression construct was made using the Web MicroRNA Designer (WMD3) and the protocol available under http://wmd3.weigelworld.org. The amiRNA precursor was cloned into pDONR221 (Invitrogen) and subsequently into pH2GW7 (Karimi *et al*., 2007b) harboring the cauliflower mosaic virus (CaMV) *35S* promoter to yield 35S:ami103. All primers used for cloning are listed in Table S1. The binary constructs were transformed into Col-0 plants by *Agrobacterium tumefaciens* (GV3101:pMP90) using the floral dip method as described by Davis, Hall, Millar, Darrah & Davis (2009). Higher order mutants with reduced expression of *ERF* genes were generated by crossing amiRNA lines with T-DNA insertion lines.

### Growth conditions, hormone and stress treatment

For hormone and stress treatments, plants were grown *in vitro* under long day (LD) conditions (16 h light/8 h dark) and 21 °C in half strength liquid Murashige and Skoog (MS) medium (for hormone treatment) or on solid MS medium (for stress treatment), in each case containing 0.1 % sucrose (Murashige & Skoog, 1962). Eleven days after germination (DAG), hormonal treatments were performed by adding the respective hormone to the liquid medium. Seedlings grown on solid medium were exposed to different stress treatments eleven DAG, including heat treatment at 42 °C in darkness, high light stress (1000 µmol m^-2^ s^-1^) instead of standard light (100‒150 µmol m^-2^ s^-1^), oxidative stress by spraying seedlings with 500 mM H_2_O_2_, drought stress by transferring seedlings to dry filter paper, or salt/osmotic stress by transplanting seedlings to MS medium including 200 mM NaCl or 200 mM mannitol, respectively, for different time periods. Control plants were treated with the respective control conditions, which were the respective mock solution in the hormone experiment, 21 °C in the heat stress experiment, standard light conditions in the high light experiment, spraying with mock solution in the oxidative stress experiment and transferring to moist filter paper in the drought experiment, or mock medium in the salt and osmotic stress experiment.

For the analysis of growth and developmental parameters, plants were grown on soil in the greenhouse under LD conditions (16 h light/8 h dark) at a light intensity of 130‒160 µmol m^-2^ s^-1^ and 21 °C. Fourteen, 21, 28, and 35 DAG rosette diameter and shoot height were determined. Furthermore, the flowering time, defined as opening of the first flower, was recorded. Leaf senescence was recorded based on visual inspection of the oldest leaves turning yellow.

For analysis of roots, plants were grown *in vitro* in vertically placed square petri dishes on half strength MS medium containing 10 g L^-1^ phytagel. The elongation of the primary root was determined from digital images between four and ten DAG using the software ImageJ (Abràmoff, Magalhaes & Ram, 2004). The number of lateral roots was determined ten DAG from the same images.

For electrolyte leakage experiments, plants were grown for two weeks under SD conditions and then for four weeks under LD conditions at 200 µmol m^-2^ s^-1^ and 20 °C during the day, 18 °C during the night (non-acclimated plants). For cold acclimation, plants were transferred to a cold chamber and cultivated under LD (90 µmol m^-2^ s^-1^) at 4 °C for additional 14 days.

### RNA analysis

Total RNA was extracted from tissues (seedlings in Fig. 2; leaves from six-week-old plants in Figure 6 and Figure S3) using the NucleoSpin RNA Plant Kit (Macherey & Nagel, Düren, Germany) according to the manufacturer’s instructions, including an on-column DNase digestion. As a control, quantitative real-time PCR (qRT-PCR) measurements using intron-specific primers for AT5G65080 were performed to confirm the absence of genomic DNA contamination (Zuther, Schulz, Childs & Hincha, 2012). For RT-PCR, 500 ng RNA were reverse transcribed using the QIAGEN OneStep RT-PCR Kit according to the manufacturer’s information (Qiagen, Hilden, Germany). The sequences of primers were as follows: *Actin2-*F, 5′-TACAACGAGCTTCGTGTTGC-3′; *Actin2-*R, 5′-GATTGATCCTCCGATCCAGA-3′; *ERF102*-F, 5′-CTGCACTTTGGTTCATCGAG-3′; *ERF102*-R, 5′-GAGATAACGGCGACAGAAGC-3′. For qRT-PCR analyses, 1 µg RNA was transcribed into cDNA by SuperScript III Reverse Transcriptase (Invitrogen) according to the manufacturer’s instructions using a combination of oligo(dT) primers and random hexamers. qRT-PCR analyses were performed as previously described by Bolt *et al*. (2017). Four biological replicates were used and each qRT-PCR experiment was performed twice. In all cases both experiments yielded similar results and one result is shown exemplarily.

### GUS staining and microscopy

Histochemical analysis to detect GUS reporter enzyme activity was performed as described by Jefferson, Kavanagh & Bevan (1987) with some modifications as described by Bolt *et al*. (2017). GUS analyses were carried out with two or three independent *pERF:GUS* lines for each of the constructs and identical expression patterns were seen. The histochemical analyses were repeated several times with plants of different age.

### Transient gene expression in *Nicotiana benthamiana* and confocal laser scanning microscopy

Subcellular localisation of GFP fused to ERF proteins was done in leaves of 6-week-old *N. benthamiana* according to Sparkes, Runions, Kearns & Hawes (2006) with the equipment described by Bolt *et al*. (2017).

### Electrolyte leakage

Electrolyte leakage was determined with detached leaves over a temperature range from -1 to -16 °C for non-acclimated plants and from -2 to -22 °C for cold acclimated plants, cooled at a rate of 4 °C h^-1^ as described in detail in Thalhammer, Hincha & Zuther (2014). Four technical replicates were analysed for each temperature point, and for each of these replicates leaves from three different plants were pooled. The temperature of 50 % electrolyte leakage (LT_50_) was calculated as the log EC50 value of sigmoidal curves fitted to the leakage values using the software GraphPad Prism3 (GraphPad Software, Inc., La Jolla, USA).

### Statistical analyses

Every experiment was conducted at least twice. Figures show data of a single experiment that is representative of two or three experiments showing similar results. Data are presented as the mean ± standard error. Statistical analyses were performed using SAS or GraphPad Instat Software (one-way ANOVA or two-way repeated measures ANOVA with Tukey’s post hoc test). Normality and homogeneity of variance were tested using the Shapiro-Wilk and Levene tests (Neter, Kutner, Nachtsheim & Wasserman, 1996). In order to meet the assumptions, data sets were transformed using log or square-root transformation. If assumptions were not met, a nonparametric Kruskal-Wallis test was carried out followed by a Mann-Whitney test to perform a pairwise comparison.

## RESULTS

### Phylogenetic analysis and description of the ERF102 to ERF105 proteins of *Arabidopsis thaliana*

According to ’The *Arabidopsis* Information Resource’ (TAIR) (Huala *et al*., 2001), *ERF102* to *ERF105* are relatively small, intronless genes with coding regions for proteins containing 300 (ERF102), 282 (ERF103), 241 (ERF104) and 221 (ERF105) amino acids. Like all AP2/ERF transcription factors they possess the characteristic AP2/ERF domain and are the only proteins in group IX with one (ERF102 and ERF103) or two (ERF104 and ERF105) putative phosphorylation sites (Nakano *et al*., 2006). Moreover, ERF102 to ERF105 possess acidic regions that might function as transcriptional activation domains (Fujimoto, Ohta, Usui, Shinshi & Ohme-Takagi, 2000). According to WoLF PSORT (Horton *et al*., 2007) ERF103 has a single nuclear localisation signal (NLS) whereas ERF102, ERF104 and ERF105 have two NLS (Figure 1a).

**Figure 1.**
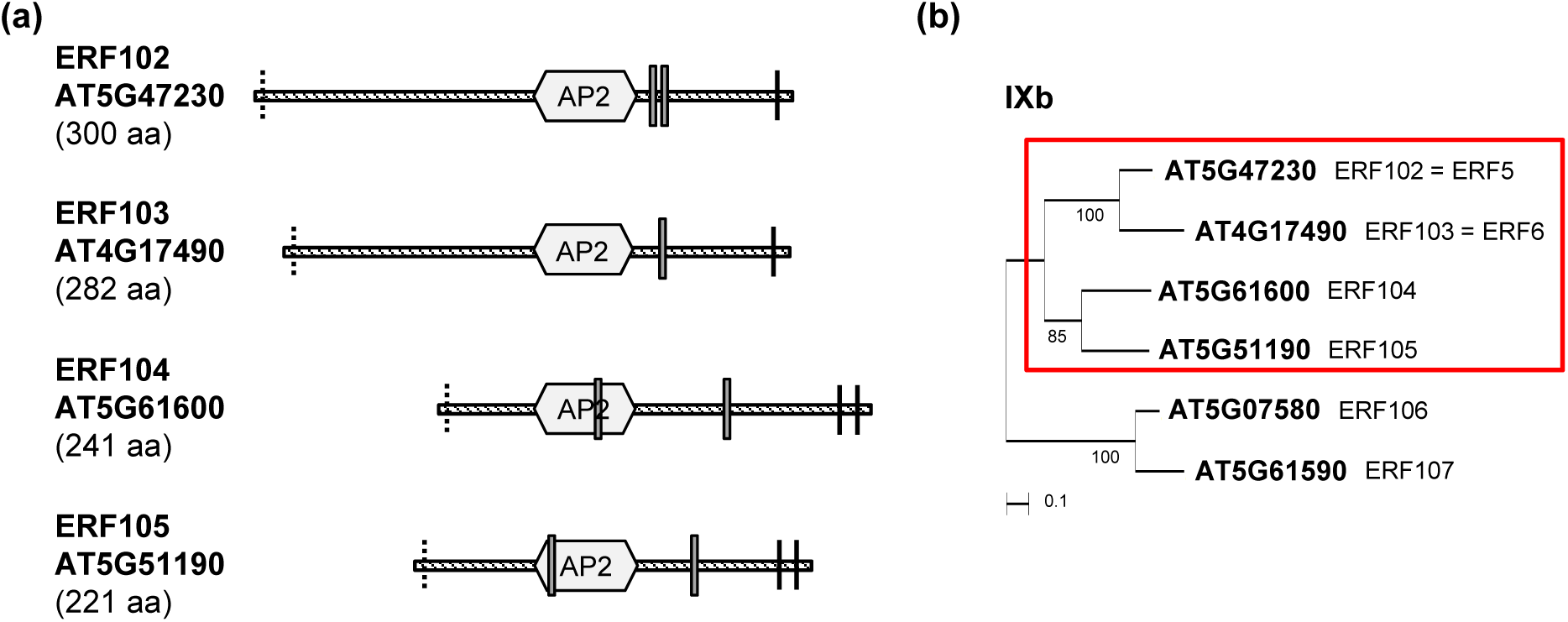
Description of the ERF102 to ERF105 proteins of *Arabidopsis thaliana*. (a) Structure of the *Arabidopsis* ERF102 to ERF105 proteins. The schematic representation shows the protein structures of ERF102 to ERF105 according to Nakano *et al*. (2006). The striped lines represent the protein sequences, the hexagons indicate the AP2/ERF DNA-binding domain, black lines putative phosphorylation sites, dashed lines the putative transactivation domains (Nakano *et al*., 2006) and grey boxes the nuclear localisation signals determined with WoLF PSORT (Horton *et al*., 2007). (b) An unrooted phylogenetic tree of group IXb ERF transcription factors showing the close evolutionary relationship between ERF102 to ERF105 (red box) that are studied. The phylogenetic tree was constructed using MEGA6, the numbers indicate bootstrap values (Tamura, Stecher, Peterson, Filpski & Kumar, 2013).

Comparison of the amino acid sequences of ERF102 to ERF105 using MUSCLE (Edgar, 2004) revealed a sequence identity of 40 % between all four proteins with high conservation of the AP2/ERF domain. The protein pairs share 67 % (ERF102 and ERF103) and 52 % (ERF104 and ERF105) amino acid identity. Phylogenetic analysis confirmed that ERF102 to ERF105 are closely related, with ERF102 and ERF103 together on one branch and ERF104 and ERF105 on the other branch of the phylogenetic tree (Figure 1b).

### The *ERF102* to *ERF105* transcription factor genes show a similar transcriptional regulation pattern

Analysis of transcriptional regulation may yield indications on functional context, therefore the previous work showing that *ERF102* to *ERF105* are regulated similarly by cold and different cold stress-related hormones, including ethylene, jasmonate and abscisic acid (Bolt *et al*., 2017), was extended. First we complemented the comparison of the hormonal transcriptional regulation of the four *ERF* genes and analysed their response to auxin and salicylic acid (SA). Auxin (NAA) rapidly and strongly induced the transcript abundances of all four *ERF* genes about 180-fold (*ERF102*), 100-fold (*ERF103*), 13-fold (*ERF104*) and 130-fold (*ERF105*) after 30 min. This increase was transient as 2 h after auxin treatment the transcript abundances were only increased between 11-fold (*ERF102*) and 2-fold (*ERF105*) (Figure 2a). In contrast, the transcript levels of all four *ERF* genes were downregulated by SA to about 50 % of the initial level after 2 h (Figure 2b).

**Figure 2.**
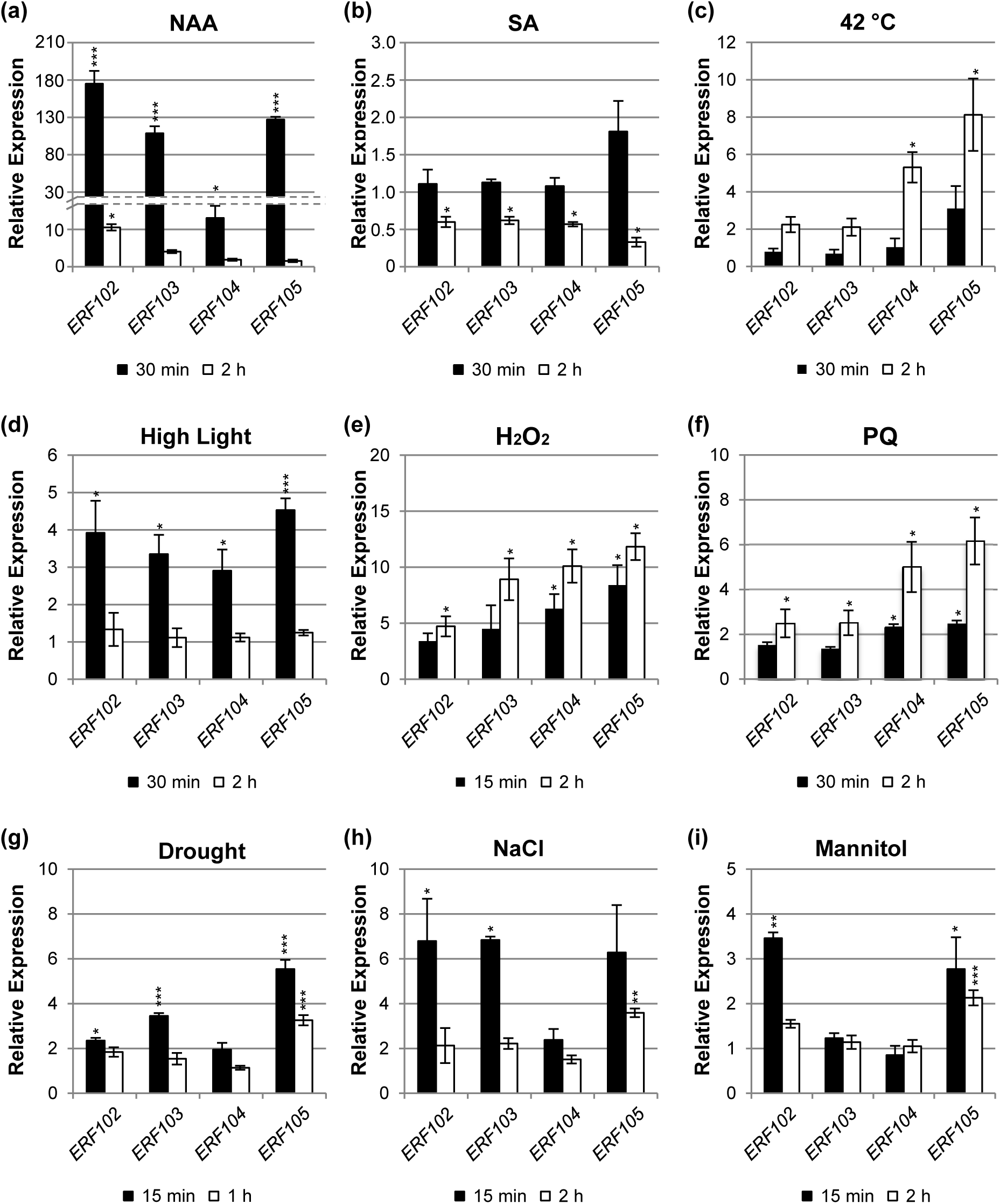
Regulation of *ERF102* to *ERF105* gene expression. Relative expression of *ERF102* to *ERF105* in eleven-day-old wild-type seedlings (eight pooled seedlings per sample) after hormone or stress treatment. (a) Auxin (10 µM NAA), (b) salicylic acid (10 mM SA), (c) heat (42 °C), (d) high light (1000 µmol m^-2^ s^-1^), (e and f) oxidative stress (e; 500 mM H_2_O_2_, f; 30 µM paraquat), (g) drought, (h) salt (200 mM NaCl) and (i) osmotic stress (200 mM mannitol). Transcript levels of wild-type samples under control conditions were set to 1 (n ≥ 4). Asterisks indicate significant differences to the respective mock treatment (*, p < 0.05; **, p < 0.01; ***, p < 0.001). Error bars represent SE.

Next, the response to different stress treatments was studied. Heat stress (42 °C) induced an upregulation of *ERF104* and *ERF105* of about 5-fold and 8-fold, respectively, after 2 h (Figure 2c). High light (1000 µmol m^-2^ s^-1^) provoked a rapid upregulation of all four genes about 4-fold (*ERF102*), 3-fold (*ERF103* and *ERF104*) and 4.5-fold (*ERF105*) after 30 min. The transcripts were back to their initial levels after 2 h (Figure 2d). Oxidative stress imposed by H_2_O_2_ treatment resulted in a rapid upregulation of all four genes after 15 min by about 3.5-fold (*ERF102*), 4.5-fold (*ERF103*), 6.5-fold (*ERF104*), and 8.5-fold (*ERF105*). After 2 h transcript levels were increased further to about 5-fold (*ERF102*), 9-fold (*ERF103*), 10-fold (*ERF104*) and 12-fold (*ERF105*) compared to the initial level (Figure 2e). Oxidative stress imposed by treatment with the superoxide-generating herbicide paraquat showed a similar result (Figure 2f). A fast transcriptional response of the *ERF* genes was also observed after drought stress that led to an about 2-fold (*ERF102* and *ERF104*), 3.5-fold (*ERF103*) and 5.5-fold (*ERF105*) upregulation of transcript levels within 15 min, which were decreased again after 1 h (Figure 2g). Salt stress (200 mM NaCl) also caused a rapid but transient upregulation of the *ERF* genes up to about 6‒7-fold for the *ERF102*, *ERF103* and *ERF105* genes (Figure 2h). Two of the genes (*ERF102*, *ERF105*) also responded rapidly to mannitol application (Figure 2i).

Taken together, the four *ERF* genes showed similar, very rapid and often transient transcriptional responses to different plant hormones, including an extraordinarily strong induction by auxin, as well as rapid, strong and often comparable responses to different stress treatments. Some individual response profiles such as stronger responses to heat by *ERF104* and *ERF105* or the lack of response to NaCl and mannitol by *ERF104* were observed as well. These partly similar stress response profiles would be consistent with overlapping functions in response to these stresses.

### *pERF102:GUS* to *pERF105:GUS* reporter genes are expressed in different tissues in Arabidopsis thaliana

Transgenic plants expressing the *GUS* reporter gene under the control of ∼2 kb of the *ERF102* to *ERF104* promoters located 5′ upstream of the coding regions were analysed to determine the tissue-specific expression of these genes.

Thirty h after imbibition, strong GUS activity of *pERF102:GUS* plants was detected in the root tip transition zone of germinated seedlings (Figure 3a) and expanded within the next 30 h within the radicle (Figure 3b). Ten DAG, *pERF102:GUS* was expressed in all root tissues except root tips and root hairs. The strongest GUS activity was observed in the vascular bundle of primary roots and in cortex cells that surround emerging lateral roots (Figure 3c–e). Weak *pERF102:GUS* expression was detected in the shoot apical meristem (SAM) of seedlings (Figure 3f).

**Figure 3.**
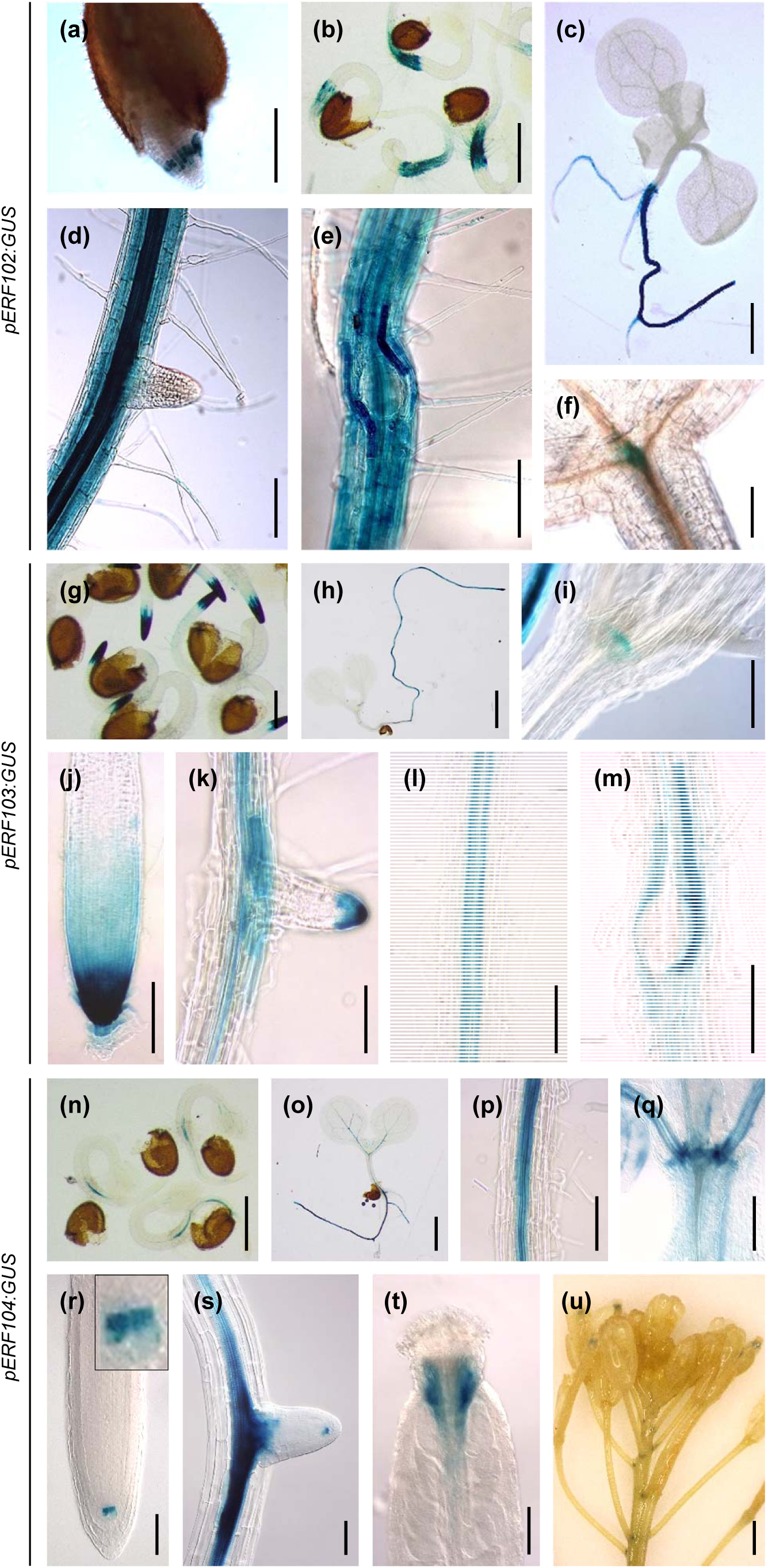
Expression of the *GUS* reporter gene under control of the *ERF102*, *ERF103* and *ERF104* promoters. Histochemical localisation of GUS activity in *Arabidopsis pERF:GUS* reporter lines. *pERF102:GUS* seedlings 30 h (a) and 60 h (b) after imbibition of seeds and ten DAG (c‒f). (a) and (b) germinating seeds, (c) whole seedling, (d) and (e) primary root with emerging lateral roots and (f) shoot apex with a stained apical meristem. *pERF103:GUS* seedlings 60 h (g) after imbibition of seeds and seven DAG (h‒m). (g) Germinating seeds, (h) whole seedling, (i) shoot apex with

*pERF103:GUS* activity was detected 60 h after imbibition in the root tip (Figure 3g) and seven DAG in the whole root (Figure 3h). Very high activity was detected in the root apical meristem (RAM) (Figure 3j). *pERF103:GUS* was also expressed in the root tip of lateral roots, but only after stage VIII of lateral root development (Péret *et al*., 2009) (Figure 3k). GUS activity was observed in the vasculature of primary roots (Figure 3l), but not in the vasculature of emerging or fully developed lateral roots, and in cortex cells that surround emerging lateral roots (Figure 3m). In shoot tissues, weak expression of *pERF103:GUS* was detected only in the shoot apex (Figure 3i).

*pERF104:GUS* expression was also detected early after germination. Sixty h after imbibition, *pERF104:GUS* was weakly expressed in the vasculature of hypocotyls and cotyledons and slightly stronger in the vasculature of radicles (Figure 3n). Seven-day-old seedlings showed GUS activity in the vascular tissues as well as in the shoot apex (Figure 3o–q). A particularly well-defined local GUS signal was noted in the quiescent center of roots (Figure 3r and 3s). In addition, GUS activity was detected in the style of the gynoecium and at the base and in the apex of siliques (Figure 3t and 3u).

As plants matured, GUS activity of *pERF102:GUS* to *pERF104:GUS* plants was present in the same tissues as in young seedlings but declined progressively (data not shown). Together, *promoter:GUS* fusions of all three *ERF* genes were predominantly expressed in root tissues, similar to *pERF105:GUS* (Bolt *et al*., 2017).

### GFP-ERF102 to GFP-ERF105 are located in the nucleus

To examine the subcellular localisation of the ERF102 to ERF104 proteins, full-length cDNAs of *ERF102* to *ERF104* were fused in frame to the 3’ end of the *GREEN FLUORESCENT PROTEIN* (*GFP*) coding sequence. The resulting *GFP-ERF102*, *GFP-ERF103* and *GFP-ERF104* fusion genes driven by the cauliflower mosaic virus (CaMV) *35S* promoter were transiently expressed in *Nicotiana benthamiana* leaf cells. Confocal imaging of GFP fluorescence in leaf cells showed that all three fusion proteins were predominantly located in the nucleus, weaker signals were derived from the cytosol (Figure 4). This pattern was similar to the predominant nuclear localisation of GFP-ERF105 (Bolt *et al*., 2017).

**Figure 4.**
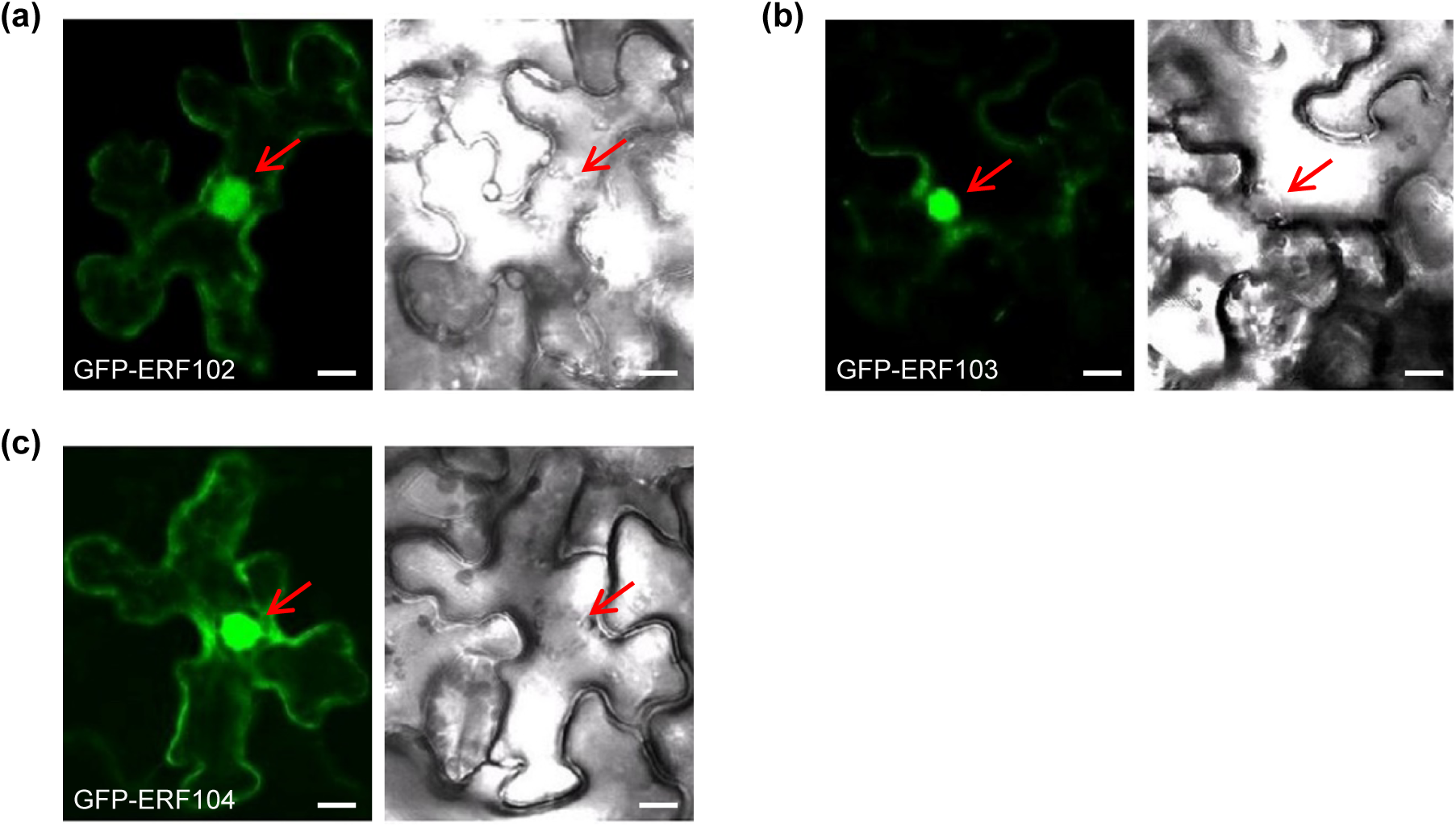
Subcellular localisation of GFP-ERF102, GFP-ERF103 and GFP-ERF104 fusion proteins. Transient expression of (a) *35S:GFP-ERF102,* (b) *35S:GFP-ERF103* and (c) *35S:GFP-ERF104* in leaf epidermis cells of *N. benthamiana* was analysed by confocal laser scanning microscopy. Left, fluorescence of GFP; right, bright field picture. The red arrows indicate the nucleus. Scale bars = 10 µm.

### Characterisation of plants with altered *ERF102* to *ERF105* expression levels

To identify and compare biological functions of the *ERF102* to *ERF104* genes, we studied transgenic lines with altered expression levels. For *ERF102*, a homozygous T-DNA insertion line (*erf102*; SAIL_46_C02) was obtained. Verification of the annotated location of the T-DNA insertion in *erf102* by sequencing revealed that the T-DNA is located at position +507 within the AP2/ERF domain (Figure S1a). RT-PCR analysis did not detect any expression of *ERF102* in *erf102* plants, suggesting that it is a null allele (Figure S1b). The morphological phenotype of the *erf102* mutant described below (Figure S2e) was fully complemented by introgression of the *35S:ERF102* gene (Figure S1c‒1f). In several available T-DNA insertion lines for *ERF103* (SALK_087356, GABI_085B06) or *ERF104* (SALK_024275, SALK_057720, SALK_152806) we detected residual *ERF* expression. Therefore, lines with a reduced *ERF103* or *ERF104* expression were constructed using artificial microRNAs (amiRNAs) (Schwab *et al*. 2006). Two independent, homozygous amiRNA expressing lines with the lowest residual expression of the target genes were selected for further experiments (Figure S2a and Bolt *et al*., 2017). Moreover, lines overexpressing *ERF102* to *ERF104* under control of the CaMV *35S* promoter were constructed and two strongly expressing lines selected (Figure S2b‒2d).

Morphological analysis of plants with reduced or increased *ERF102* to *ERF104* expression revealed in most cases only slight differences of shoot growth compared to wild-type plants. Furthermore, plants with altered expression of *ERF102*, *ERF103* or *ERF104* flowered at the same time as wild-type plants and showed a similar onset of leaf senescence (data not shown). In contrast, root elongation, the formation of lateral roots as well as the lateral root density was more strongly affected by altered expression of these genes (Figure 5c–5e).

**Figure 5.**
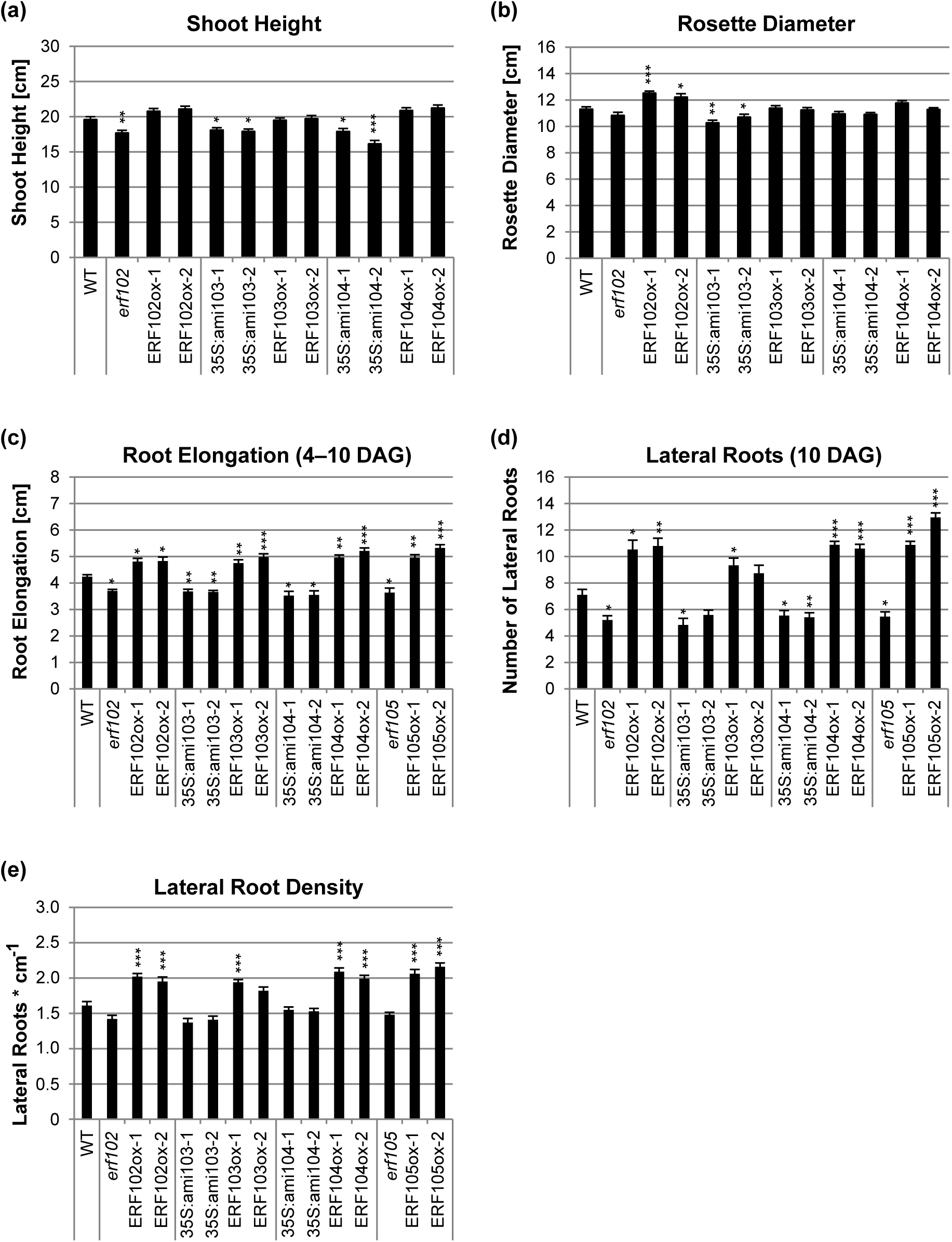
Shoot and root growth of lines with altered *ERF102* to *ERF105* expression levels. Shoot height (a) and rosette diameter (b) of 35-day-old plants grown on soil. (c) Elongation of the primary root determined between four and ten DAG (c), number of lateral roots (d) and lateral root density (e) determined ten DAG of plants grown on half-strength MS medium. Asterisks indicate significant differences to the wild type (n ≥ 30), (*, p < 0.05; **, p < 0.01; ***, p < 0.001). Error bars represent SE.

The *erf102* mutant exhibited an about 10 % reduced shoot height compared to the wild type. Overexpressing lines of *ERF102* exhibited a slightly but not significantly increased shoot height as well as a 10 % (ERF102ox-1) and 8 % (ERF102ox-2) bigger rosette diameter (Figure 5a and 5b). Moreover, ten DAG *erf102* exhibited 27 % less and ERF102ox-1 and ERF102ox-2 48 % and 51 % more lateral roots compared to wild type (Figure 5d). Lateral root density was increased 29‒31 % in the ERF102ox lines (Figure 5e).

Both 35S:ami103 lines were smaller in size, with an 8 % reduced shoot height and a 6‒ 9 % reduced rosette diameter compared to the wild type, while *ERF103* overexpression did not cause phenotypic differences in shoot height and rosette size (Figure 5a and 5b). Primary root elongation was about 13 % lower in both 35S:ami103 lines whereas ERF103ox-1 and ERF103ox-2 exhibited 12 % and 17 % longer primary roots compared to wild type (Figure 5c). Similarly, 35S:ami103 lines had up to 32 % less and ERF103ox plants up to 31 % more lateral roots than wild type (Figure 5d).

35S:ami104 lines had a 9 % (35S:ami104-1) and 18 % (35S:ami104-2) reduced shoot height, but an unchanged rosette diameter (Figure 5a and 5b). Primary root elongation of 35S:ami104 lines was slightly reduced (about 13 % in 35S:ami104-2) and enhanced by up to 29 % in *ERF104* overexpressing lines (Figure 5c). The number of lateral roots was reduced by about 20 % in both 35S:ami104 lines, while ERF104ox-1 and ERF104ox-2 exhibited 57 % and 53 % more lateral roots (Figure 5d) and had a 30 % and 22 % higher lateral root density compared to wild type (Figure 5e).

Bolt *et al*. (2017) described that the shoot phenotype of *erf105* and ERF105ox lines resembled the wild type. Here, root analysis revealed 23 % less lateral roots in the *erf105* mutant compared to wild type (Figure 5c). ERF105ox lines showed a 17-25 % higher primary root elongation, 53-83 % more lateral roots and a 31-44 % higher lateral root density compared to wild type (Figure 5c–5e).

To examine a potentially redundant role of the four *ERF* genes, several higher order mutants were generated, namely *erf102* 35S:amiERF103, *erf102* 35S:amiERF104, *erf105* 35S:amiERF103, and *erf102* 35S:amiERF104/105. These lines include all possible combinations of at least two *ERF* genes that are mutated or have a lowered expression, except combined loss of function of *ERF103* and *ERF104*. Higher order mutants did not show a phenotypic additive effect compared to the respective single mutants with respect to rosette diameter, shoot height, primary root elongation, number of lateral roots and flowering time (data not shown). These results suggest that *ERF102* to *ERF105* are not acting redundantly on growth regulation. However, we cannot exclude that the degree of downregulation achieved by amiRNAs is insufficient to uncover redundant gene activities.

### Analysis of the functional redundancy of the *ERF102* to *ERF105* genes in the cold acclimation response

ERF105 is a positive regulator of *Arabidopsis* freezing tolerance and cold acclimation (Bolt *et al*., 2017). Therefore, we analysed whether the *ERF102* to *ERF104* genes, which are also regulated by cold (Bolt *et al*., 2017; Lee *et al*., 2005; Park *et al*., 2015; Vogel *et al*., 2005), also play a role in regulating freezing tolerance and cold acclimation. To this end, we studied the transcript accumulation of selected cold responsive genes in *ERF* single and double mutants and analysed the freezing tolerance of these mutants.

First, we examined the expression levels of selected cold-responsive genes in plants with reduced or enhanced expression of a single *ERF102* to *ERF104* gene before (non-acclimated, NA) and after 14 d of cold acclimation (ACC14) and compared these to wild type. The transcript levels of cold-responsive genes were in all lines similar to wild type (Figure S3), which contrasts with the strongly altered transcript levels displayed by the *erf105* mutant and *ERF105* overexpressing lines (Bolt *et al*., 2017).

The analysis of higher order mutants revealed that under non-acclimated (NA) conditions the steady state mRNA levels of *CBF1*, *CBF2*, *COR15A*, and *COR15B* were up to 60 % lower in the *erf105* 35S:ami103-1 plants compared to those of the wild type (Figure 6). In all other mutant combinations the basic expression level of these cold-responsive genes was slightly, but not significantly lower than in the wild type. After 14 d of acclimation at 4 °C (ACC14), the expression levels of these genes were elevated between 2- and 5-fold in wild type compared to NA plants. ACC14 plants with mutated *ERF102* or *ERF105* genes combined with reduced expression of *ERF103* or *ERF104* showed, in most cases, a lower induction of the cold-responsive genes. For example, the induction levels of *CBF2* and *COR15B* were reduced in all hybrid lines to about 50 % of the wild-type level. Strikingly, the induction of *CFB3* was completely absent in all mutant lines while it was induced about 2-fold in wild type. In contrast, *ZAT12* gene expression showed a stronger increase in *erf102* 35S:ami103-1, *erf102* 35S:ami104-2 and *erf105* 35S:ami103-1 than in wild type (Figure 6f).

**Figure 6.**
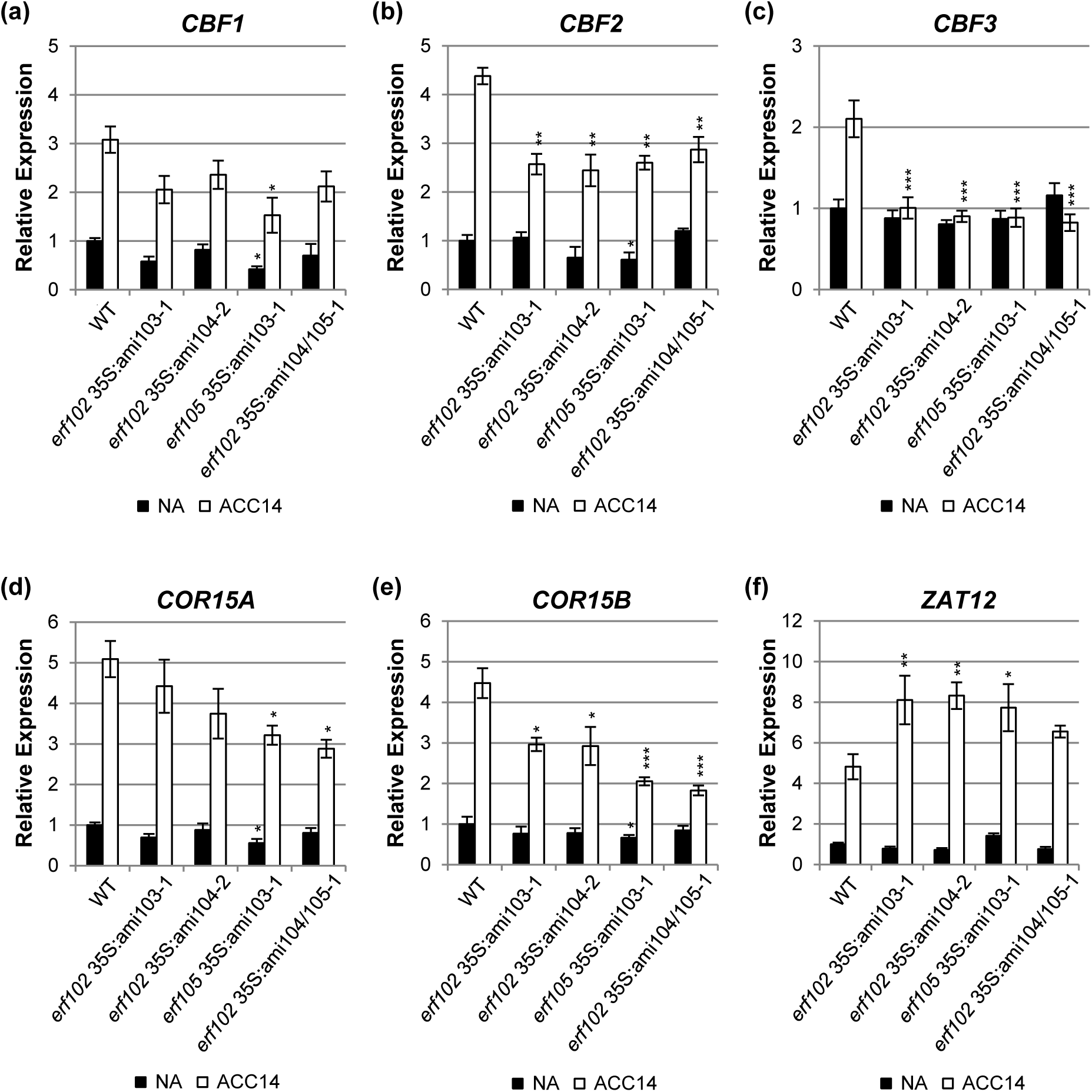
Expression of selected cold-responsive genes in lines with reduced *ERF102* to *ERF105* expression. Relative expression of *CBF1* (a), *CBF2* (b), *CBF3* (c), *COR15A* (d), *COR15B* (e) and *ZAT12* (f) genes in lines with reduced *ERF102* to *ERF105* expression before (non-acclimated, NA) and after 14 days (acclimated, ACC14) of cold acclimation at 4 °C. Transcript levels of wild-type samples under non-acclimated conditions were set to 1 (n ≥ 4). Asterisks indicate significant differences to the respective wild-type condition (*, p < 0.05; **, p < 0.01; ***, p < 0.001). Error bars represent SE.

Next, we determined the freezing tolerance of plants with reduced *ERF102, ERF103* and *ERF104* gene expression before and after 14 d of cold acclimation at 4 °C by an electrolyte leakage assay of detached leaves (Thalhammer *et al*., 2014). To take into account the almost complete arrest of plant growth at 4 °C, the electrolyte leakage assay was performed at the same developmental state for both NA and ACC plants. *erf105* mutant plants used as positive control showed higher LT_50_ (temperature of 50 % electrolyte leakage) values (−3.99 ± 0.13 °C in NA plants and −8.99 ± 0.17 °C in ACC14 plants) compared to wild type (−4.7 ± 0.11 °C in NA plants and −10.82 ± 0.12 °C in ACC14 plants) (Figure 7a), which is consistent with previous results (Bolt *et al*., 2017). In contrast, *erf102*, 35S:ami103-1 and 35S:ami104-2 plants did not show differences in LT_50_ values compared to wild type. Also, overexpression of single *ERF102*, *ERF103* or *ERF104* genes did not lead to altered freezing tolerance under NA conditions (Figure S4). The behavior of the overexpressing lines in response to acclimation was not tested. Analysis of the freezing tolerance of higher order mutants revealed that only the *erf105* 35S:ami103-1 plants showed higher LT_50_ values (−4.93 ± 0.12 °C) compared to wild type (−5.46 ± 0.12 °C) under NA conditions (Figure 7b). Following cold acclimation, several combinations exhibited higher LT_50_ values compared to wild type (−9.54 ± 0.18 °C). The strongest change was shown by *erf102* 35S:ami103-1 (−7.89 ± 0.24 °C), while *erf105* 35S:ami103-1 (−8.78 ± 0.25 °C) as well as *erf102* 35S:ami104/105-1 (−8.79 ± 0.25 °C) showed smaller effects. In contrast, *erf102* 35S:ami104-2 showed a similar LT_50_ as wild type after cold acclimation (Figure 7b).

**Figure 7.**
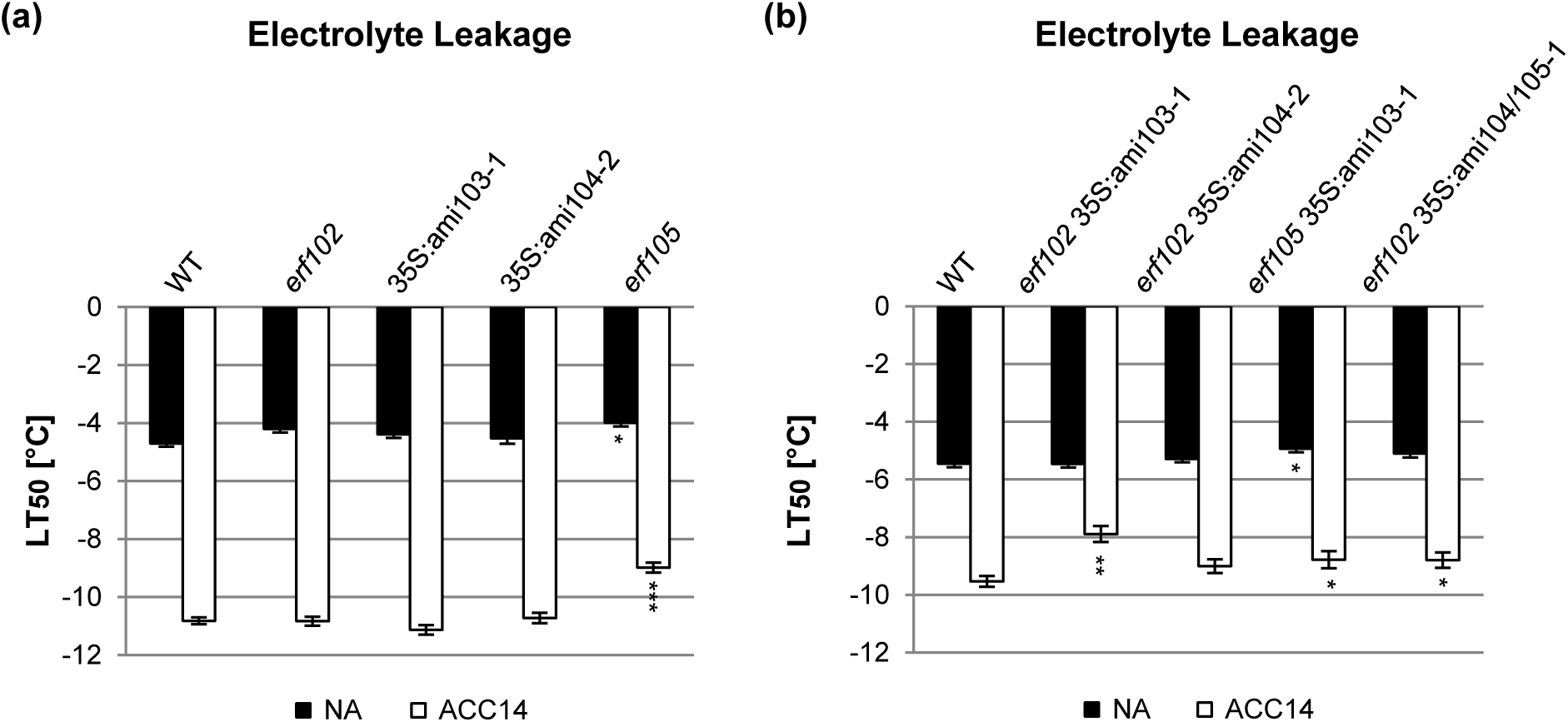
Electrolyte leakage assays of lines with reduced *ERF102* to *ERF105* expression. Electrolyte leakage assays on detached leaves of lines with mutations or reduced expression affecting single *ERF* genes (a) or several *ERF* genes (b) before (non-acclimated, NA) and after 14 days (acclimated, ACC14) of cold acclimation at 4 °C. The bars represent the means ± SE from four replicate measurements where each replicate comprised leaves from three plants. Asterisks indicate significant differences to the wild type (*, p < 0.05; **, p < 0.01; ***, p < 0.001).

## DISCUSSION

Recently, we reported that *ERF102* to *ERF105* are regulated by cold and different cold stress-related hormones, and we demonstrated that *ERF105* has a function in the freezing tolerance and cold acclimation of *Arabidopsis* (Bolt *et al*., 2017). In the present study we significantly extended this work and first investigated further expression characteristics of the gene family members and then explored their potentially redundant roles in regulating plant growth and the cold acclimation response.

### *The* ERF102 *to* ERF105 *genes show overlapping expression patterns*

The similar profiles of gene expression in response to hormone or stress treatment are consistent with a partial functional redundancy of *ERF102* to *ERF105*. For instance, all genes were rapidly downregulated by SA (Figure 2b) and upregulated by high light or H_2_O_2_ (Figure 2e and 2f). Network analysis of publicly available transcriptome data using for instance GeneMANIA (Warde-Farley *et al*., 2010) also showed that these four *ERF* genes are co-regulated and co-expressed in a large number of conditions including numerous hormone and chemical treatments (Figure S5). However, some individual response profiles were discovered as well. Thus, not all four *ERF* genes were transcriptionally regulated by heat, drought, NaCl, or mannitol (Figure 2). Together, the analysis of transcriptional regulation is in line with the idea that *ERF102* to *ERF105* have roles in multiple hormone and stress responses as was shown for these and other ERFs in a number of cases (Bethke *et al*., 2009; Dubois *et al*., 2013, 2015; reviewed by Licausi, Ohme-Takagi & Perata 2013; Mase *et al*., 2013; Meng *et al*., 2013; Moffat *et al*., 2012; Moore *et al*., 2014; Sewelam *et al*., 2013; Son *et al*., 2012; Vogel *et al*., 2014; Xie *et al*., 2019).

### The ERF102 to ERF105 genes have a limited impact on plant growth

The tissue-specific expression patterns of *pERF102:GUS* to *pERF105:GUS* are partly overlapping, which is in accordance with a redundant function of the ERF proteins. All four genes are predominantly expressed in the root, only for *pERF105:GUS* a significant expression was detected also in several shoot tissues such as vasculature, apical shoot and stomata (Bolt *et al*., 2017). Expression of all four *pERF-GUS* reporter genes was visible shortly after germination in different cell types of the radicle and later in distinct root tissues and cell types. For example, *pERF102:GUS*, *pERF103:GUS* and *pERF105:GUS* were expressed in the cortex cells that surround emerging lateral roots. Interestingly, expression of *ERF102*, *ERF103* and *ERF105* is regulated by cytokinin and auxin, two key hormones of lateral root development (Benková *et al*., 2003; Casimiro *et al*., 2003; Chang, Ramireddy & Schmülling, 2013, 2015; Swarup *et al*., 2008). However, insertional mutants or amiRNA lines did not reveal a major role of these genes in regulating root architecture. 35S:ami103 and 35S:ami104 lines had shorter roots and most loss-of-function mutants formed less lateral roots. However, the differences were small and the lateral root density mostly not significantly altered (Figure 5c-e). Opposite and stronger phenotypic changes were noted in the respective overexpressing lines, which had longer roots, an increased number (by up to ∼85 %) of lateral roots and a higher lateral root density. Although overexpression experiments may produce artefacts and are not fully conclusive they have been often informative about the functional context of a given gene. Loss-of-function phenotypes of genes regulating root architecture can be subtle or depend on the environmental or developmental context (Motte, Vanneste & Beeckman, 2019) and thus might have gone unnoticed in the *erf* mutants. The strong regulation of the four *ERF* genes by different stressors suggests that they might be particularly relevant under stressful conditions. It cannot be excluded that members of the *ERF105* gene subfamily studied here contribute to regulating root architecture under specific environmental conditions, this requires further investigation.

Among the expression sites of the four *ERF* genes, the expression of *pERF104:GUS* in the quiescent center (Figure 3r) particularly intriguing. Noteworthy, among the direct targets of ERF104 is the transcription factor gene *SCARECROW* (*SCR*) (Sparks *et al*., 2016). SCR is, together with SHORTROOT, essential for quiescent center specification and maintenance (Salvi *et al*., 2018; reviewed by Benfey, 2016). Further, in a yeast two-hybrid screen the transcription factor MYB56/BRASSINOSTEROIDS AT VASCULAR AND ORGANIZING CENTER (BRAVO) was identified as an interactor of ERF104 (our unpublished result). MYB56/BRAVO represses cell divisions in the quiescent center thus counteracting SCR (Di Laurenzio *et al*., 1996; Vilarrasa-Blasi *et al*., 2014). It is known that interaction with other transcription factors modulates the activity of ERFs (Licausi *et al*., 2013; Xie *et al*., 2019). While these data suggest that ERF104 might be part of the transcription factor network in the quiescent center, we have been unable to detect any changes of cellular organisation in the quiescent center and surrounding cells nor did we detect altered *SCR* gene expression in the 35S:ami104 and ERF104ox lines (data not shown). It could be that the decrease in *ERF104* expression obtained in the amiRNA lines is not sufficient to cause a strong loss-of-function phenotype, analysis of a null mutation could be more informative.

### The ERF102 to ERF105 genes redundantly regulate the response to cold stress

One important goal of this work was to analyse the possible roles of the ERF105-related transcription factors in the response to cold stress. *ERF102* to *ERF105* are rapidly cold-induced (Bolt *et al*., 2017) in parallel with the first wave transcription factors of the cold stress response including the *CBF* genes (Park *et al*., 2015). Mutation or reduced expression of *ERF102, ERF103* or *ERF104* single genes did not lead to an altered freezing tolerance. In case of the amiRNA lines this could be due to residual gene expression (Figure 7a and S1). Thus, among the four genes only the mutation of *ERF105* resulted in a decreased freezing tolerance before and after cold acclimation compared to wild type underpinning its primary role (Figure 7a and Bolt *et al*., 2017). However, the analysis of freezing tolerance of higher order mutants indicated that *ERF102* and *ERF103* also play a role in cold acclimation, since the reduced expression of both genes resulted in altered expression of cold response genes (Figure 6) and higher freezing sensitivity (Figure 7b). The eventual role of ERF104 cannot be determined with certainty as only amiRNA lines were available and not all combinations with other *ERF* genes were tested. 35S:ami104 lines in combination with the *erf102* mutation showed an altered expression of cold-responsive genes similar to other double mutant combinations (Figure 6) and the LT_50_ value was higher than in wild type although the significance was below the threshold (p < 0.05), indicating that ERF104 might be involved in the response to cold as well. Our attempts to demonstrate a role of these *ERF* genes at low temperatures in the root as was reported for *CRF2* and *CRF3* belonging to a different class of *ERF* genes (Jeon et al., 2016), have failed. Such an activity could, as was stated above, be masked by incomplete loss of function and/or the unknown nature of their specific activities.

Based on transcript data which show a lowered activation of *CBF* and *COR* genes in *erf* gene mutants after cold acclimation (Figure 6), ERF102, ERF103 and ERF104 may also play a role upstream of these genes as was suggested for ERF105 (Bolt *et al*., 2017). Increased *CBF3* expression upon cold acclimation was even completely lacking in the *erf* mutants (Figure 6c) but the gene was still cold responsive at earlier time points although with a reduced amplitude as compared to wild type (Figure S6). A proximity of the four *ERF* genes to the CBF regulon was also suggested by the result of the network analysis which placed several proteins that are part of the CBF regulon (CBF2/DREB1c, ZAT10 und RAP2.13/RAP2.4) in the vicinity of ERF102 to ERF105 (Figure S5).

The lower activation of the *CBF* and *COR* genes in cold-acclimated *erf* gene mutants could be at least partially due to enhanced expression of another gene belonging to the CBF regulon, *ZAT12* (Figure 6f). *ZAT12* encodes a zinc-finger protein known to be a negative regulator of the CBF regulon and is usually induced in parallel with *CBF* and *COR* genes providing a negative regulatory feedback loop (Vogel *et al*., 2005). The higher expression of *ZAT12* in the *erf* higher order mutants suggests that these *ERF* genes may act as negative regulators of *ZAT12* expression and in this way as positive regulators of *CBF* and *COR* genes. Notably, the *ZAT12* gene does not possess the specific DNA-binding motif of ERF transcription factors, the GCC-box, in its promoter region (Hao *et al*., 1998) suggesting that additional factors might be required for its repression by ERFs.

Knockout/knockdown of single *ERF102* to *ERF104* genes did not cause an altered transcript level of cold-responsive genes after 14 d of cold acclimation (Figure S3), which is again in line with the assumption that these *ERF* genes may have redundant roles. Lines overexpressing *ERF102* to *ERF104* did neither show a differential expression of cold-responsive genes nor an altered freezing tolerance (Figure S3 and S4), similar to *ERF105* overexpressing lines (Bolt *et al*., 2017). It is possible that ERF102 to ERF105 are required for the transcriptional activation of these target genes but are not the rate-limiting factors, for example because they function as part of a complex. Alternatively, activity of these proteins under cold may depend on additional regulatory steps such as phosphorylation which could be transient. Indeed, the phosphorylation of ERF102 to ERF104 by MPK3 and/or MPK6 was shown (Bethke *et al*., 2009; Son *et al*., 2012; Wang, Du, Zhao, Miao & Song, 2013) and functions of MPK3 and MPK6 in the cold signalling pathway have been described (Kim *et al*., 2017; Li *et al*., 2017; Zhao *et al*., 2017).

Taken together, the data document redundant functions of *ERF102* to *ERF105* in response to cold. Notably, combined action of related *ERF* transcription factor genes has also been reported in other cases (Jeon, Cho, Lee, Van Binh & Kim, 2016; Kim, Jang & Park, 2016). Future work will investigate how the ERF102 to ERF105 proteins are integrated in the extensive transcriptional network governing the response to cold.

## SUPPORTING INFORMATION

**Figure S1. Characterisation of the *erf102* mutant SAIL_46_C02.** (a) Structure of the *Arabidopsis ERF102* (AT5G47230) gene. The black line denotes the untranslated region, the black box represents the exon, the T-DNA insertion at position +507 is shown by a triangle. The positions of primers that were used for RT-PCR are indicated by arrows. (b) RT-PCR analysis of *ERF102* expression using total RNA extracted from seedlings of wild type and *erf102*. The *Actin2* gene was used as internal control. (c‒f) Complementation of the *erf102* mutant by introgression of the *35S:ERF102* gene. Shoot height (c) and rosette diameter (d) of 35-day-old plants. (e) Elongation of the primary root and (f) number of lateral roots of plants grown on half-strength MS medium. Asterisks indicate significant differences to the wild type (n ≥ 30), (*, p < 0.05; **, p < 0.01). Error bars represent SE.

**Figure S2. Analysis of lines with altered *ERF102* to *ERF104* expression levels.** (a‒d) Relative expression level of *ERF* genes in eight pooled eleven-day-old seedlings of wild type, lines expressing amiRNA directed against *ERF103* (a) and lines overexpressing *ERF102* (b), *ERF103* (c,) or *ERF104* (d). Transcript levels of wild-type samples were set to 1 (n ≥ 4). Asterisks indicate significant differences to the wild type (***, p < 0.001). Error bars represent SE. (e‒g) Shoot phenotype of plants grown 35 days under long day conditions. The pictures are complementary to the data shown in Figure 5a and 5b.

**Figure S3. Expression of selected cold-responsive genes in lines with reduced or enhanced *ERF102* to *ERF104* expression.** Relative expression of *CBF1* (a), *CBF2* (b), *COR15A* (c), and *COR15B* (d) genes in lines with reduced or enhanced *ERF102* to *ERF104* expression before (non-acclimated, NA) and after 14 days (acclimated, ACC14) of cold acclimation at 4 °C. Transcript levels of wild-type samples under non-acclimated conditions were set to 1 (n ≥ 4). Error bars represent SE.

**Figure S4. Electrolyte leakage assays of lines with enhanced *ERF102* to *ERF104* expression.** Electrolyte leakage assays on detached leaves of lines overexpressing *ERF102*, *ERF103* or *ERF104* before (non-acclimated, NA) and after 14 days (acclimated, ACC14) of cold acclimation at 4 °C. The bars represent the means ± SE from four replicate measurements where each replicate comprised leaves from three plants.

**Figure S5. Network of co-localisation, co-expression, genetic and physical interactions of ERF105.** The blue connecting lines between two genes represent co-localisation, purple lines co-expression, green lines genetic interactions and red lines physical interactions. ABI1, ABA INSENSITIVE 1; AZF3, ZINC-FINGER PROTEIN 3; CAF1-9, CCR4-ASSOCIATED FACTOR 1 HOMOLOG 9; CYP707A3, CYTOCHROME P450, FAMILY 707, SUBFAMILY A, POLYPEPTIDE 3; DREB1C (CBF2), DEHYDRATION-RESPONSE ELEMENT-BINDING PROTEIN 1C/C-REPEAT-BINDING FACTOR 2; ERF, ETHYLENE RESPONSE FACTOR; PP2CA, PROTEIN PHOSPHATASE 2CA; PUMP4, PLANT UNCOUPLING MITOCHONDRIAL PROTEIN 4; RAP2-13 (RAP2.4/WIND), RELATED TO AP2 13; SZF1, SALT-INDUCIBLE ZINC-FINGER; ZAT10 (STZ), ZINC FINGER PROTEIN 10 (SALT TOLERANCE ZINC FINGER). Analysis was done using GeneMANIA (Warde-Farley *et al*., 2010).

**Figure S6. Expression of selected cold-responsive genes in lines with reduced *ERF102* to *ERF105* expression.** Relative expression of *CBF1*, *CBF2* and *CFB3* genes in lines with reduced *ERF102* to *ERF105* after 4 h of cold treatment at 4 °C. Transcript levels of wild-type samples under control conditions were set to 1 (n ≥ 4). Asterisks indicate significant differences to the wild type (*, p < 0.05; ***, p < 0.001). Error bars represent ± SE.

**Table S1. Sequences of primers used for cloning.** Small letters in the primer sequences indicate the integrated *attB4*- or *attB1*-sites for cloning DNA fragments into the vector pDONR P4-P1R. Small italic letters in the primer sequences indicate the integrated *attB1*- or *attB2*-sites for cloning DNA fragments into the vector pDONR221. Underlined letters are the nucleotides added to keep the sequence in the right frame.

## ACKNOWLEDGEMENTS

We acknowledge funding by Deutsche Forschungsgemeinschaft (Collaborative Research Centre 973, www.sfb.973).

