## Supplemental data for "Characterisation of the *ERF102* to *ERF105* genes of *Arabidopsis thaliana* and their role in the response to cold stress"

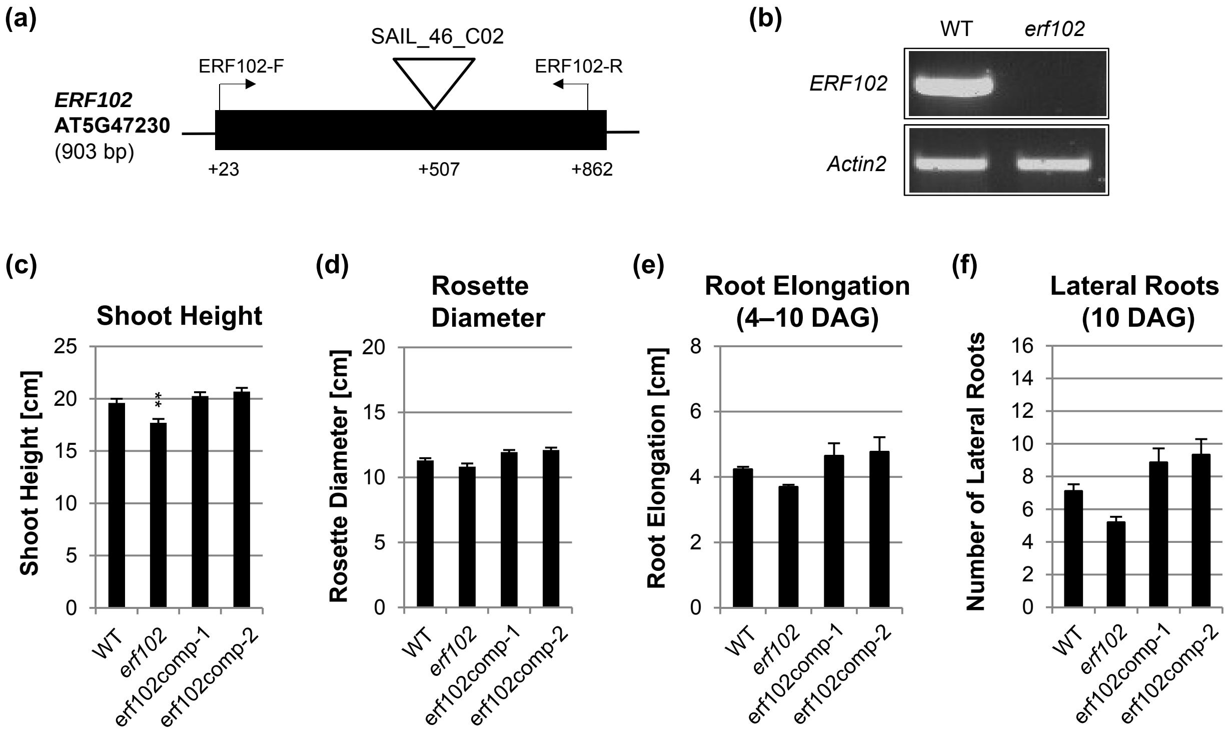


**Figure S1. Characterisation of the *erf102* mutant SAIL_46_C02.** (a) Structure of the *Arabidopsis ERF102* (AT5G47230) gene. The black line denotes the untranslated region, the black box represents the exon, the T-DNA insertion at position +507 is shown by a triangle. The positions of primers that were used for RT-PCR are indicated by arrows. (b) RT-PCR analysis of *ERF102* expression using total RNA extracted from seedlings of wild type and *erf102*. The *Actin2* gene was used as internal control. (c–f) Complementation of the *erf102* mutant by introgression of the *35S:ERF102* gene. Shoot height (c) and rosette diameter (d) of 35-day-old plants. (e) Elongation of the primary root and (f) number of lateral roots of plants grown on half-strength MS medium. Asterisks indicate significant differences to the wild type ($n \geq 30$), (*, $p < 0.05$; **, $p < 0.01$). Error bars represent SE.

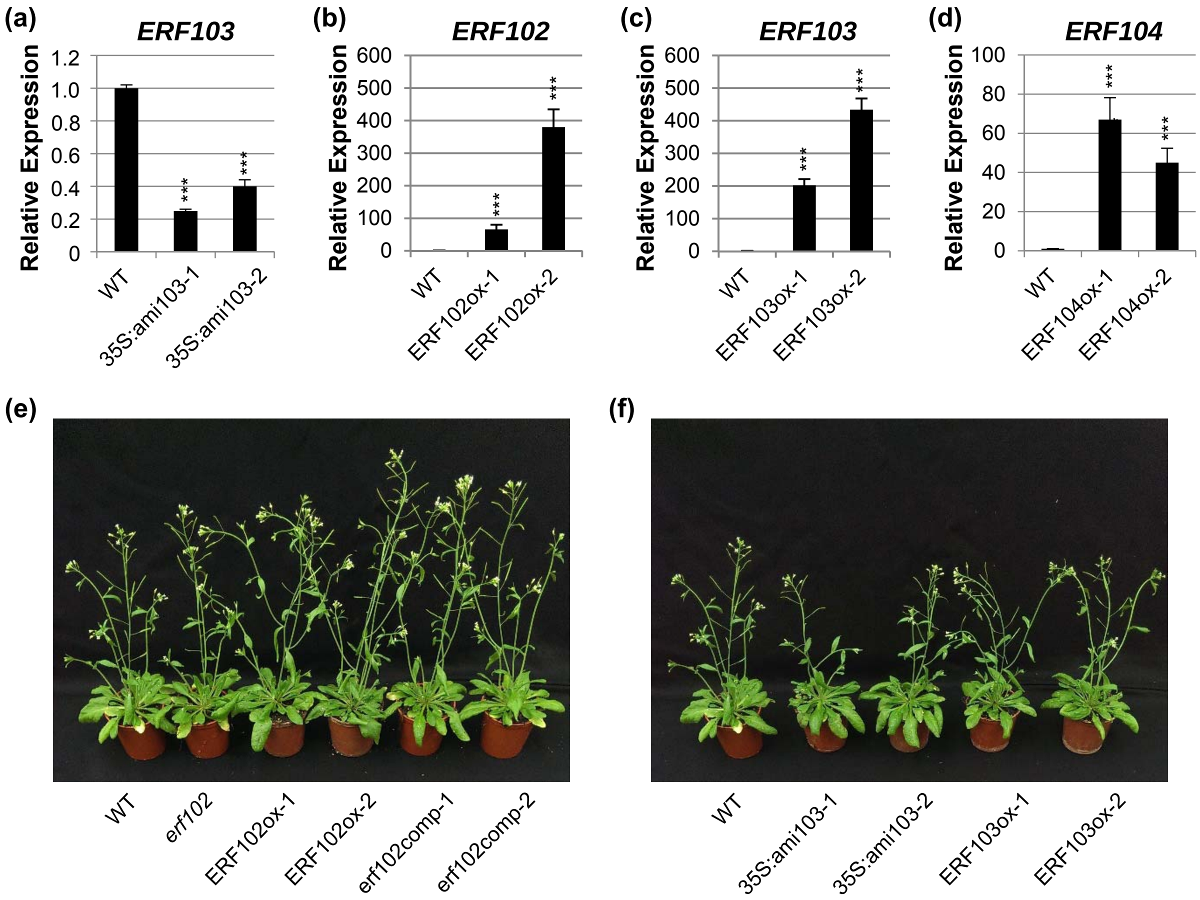


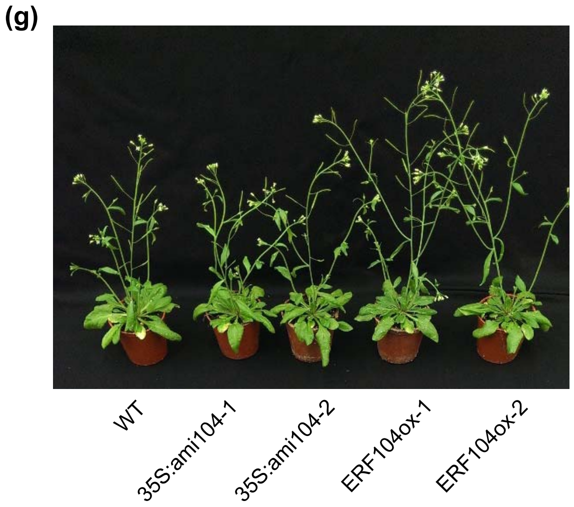


**Figure S2. Analysis of lines with altered *ERF102* to *ERF104* expression levels.** (a–d) Relative expression level of *ERF* genes in eight pooled eleven-day-old seedlings of wild type, lines expressing amiRNA directed against *ERF103* (a) and lines overexpressing *ERF102* (b), *ERF103* (c) or *ERF104* (d). Transcript levels of wild-type samples were set to 1 ($n \geq 4$). Asterisks indicate significant differences to the wild type (***, $p < 0.001$). Error bars represent SE. (e–g) Shoot phenotype of plants grown 35 days under long day conditions. The pictures are complementary to the data shown in Figure 5a and 5b.

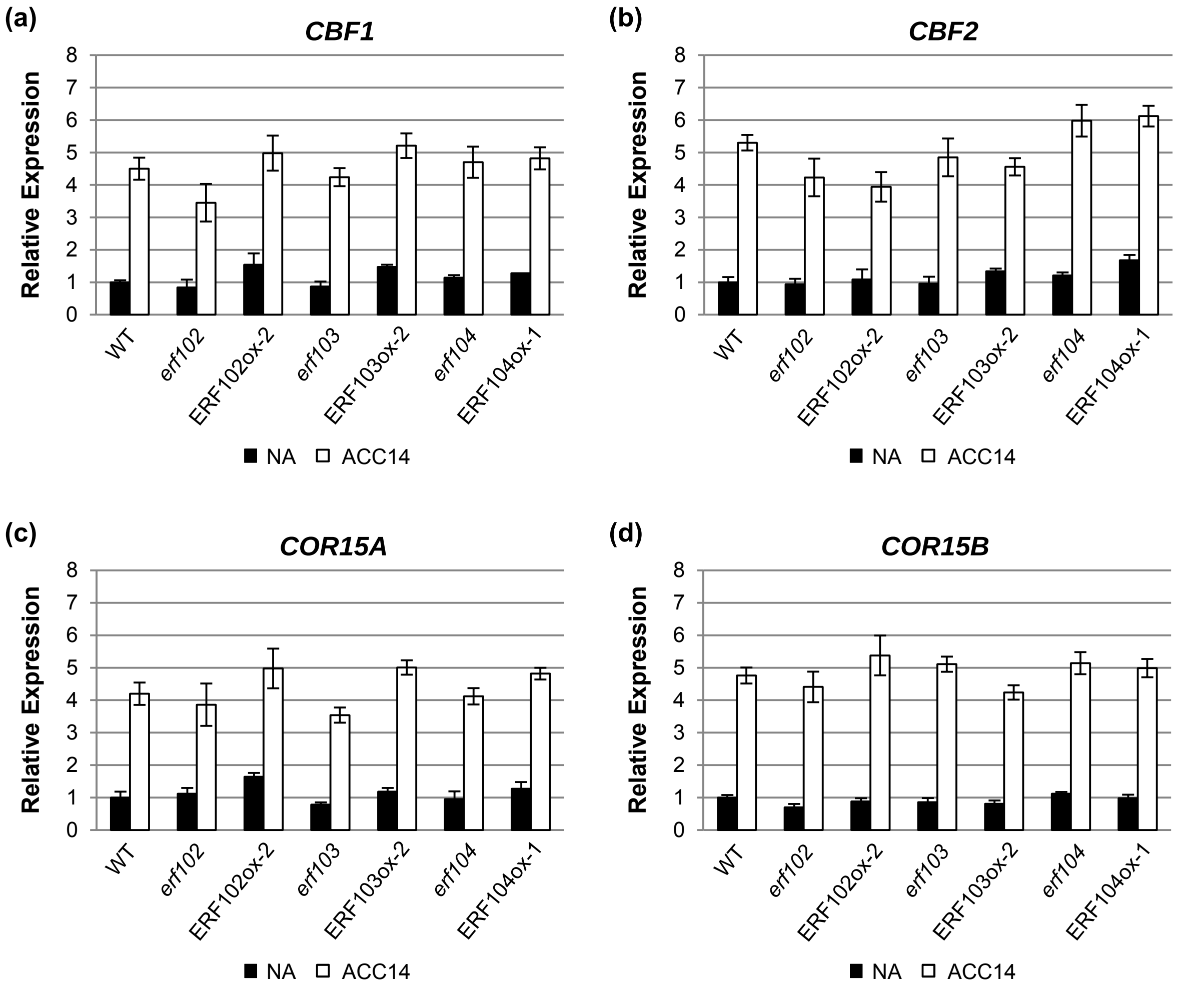


**Figure S3. Expression of selected cold-responsive genes in lines with reduced or enhanced *ERF102* to *ERF104* expression.** Relative expression of *CBF1* (a), *CBF2* (b), *COR15A* (c), and *COR15B* (d) genes in lines with reduced or enhanced *ERF102* to *ERF104* expression before (non-acclimated, NA) and after 14 days (acclimated, ACC14) of cold acclimation at 4 °C. Transcript levels of wild-type samples under non-acclimated conditions were set to 1 ($n \geq 4$). Error bars represent SE.

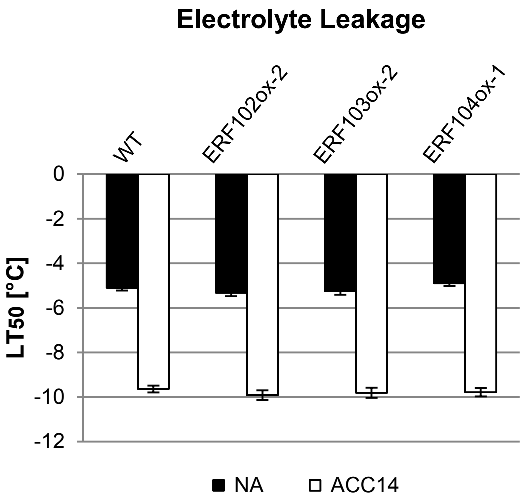


**Figure S4. Electrolyte leakage assays of lines with enhanced *ERF102* to *ERF104* expression.** Electrolyte leakage assays on detached leaves of lines overexpressing *ERF102*, *ERF103* or *ERF104* before (non-acclimated, NA) and after 14 days (acclimated, ACC14) of cold acclimation at 4 °C. The bars represent the means ± SE from four replicate measurements where each replicate comprised leaves from three plants.

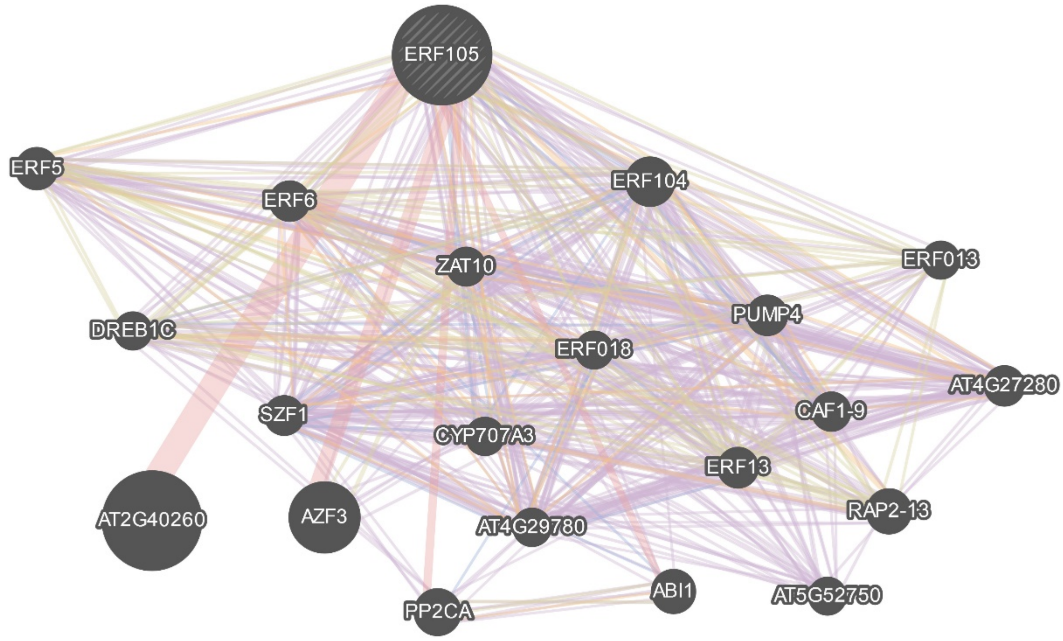


**Figure S5. Network of co-localisation, co-expression, genetic and physical interactions of ERF105.** The blue connecting lines between two genes represent co-localisation, purple lines co-expression, green lines genetic interactions and red lines physical interactions. ABI1 = ABA INSENSITIVE 1; AZF3 = ZINC-FINGER PROTEIN 3; CAF1-9 = CCR4-ASSOCIATED FACTOR 1 HOMOLOG 9; CYP707A3 = CYTOCHROME P450, FAMILY 707, SUBFAMILY A, POLYPEPTIDE 3; DREB1C (CBF2) = DEHYDRATION-RESPONSE ELEMENT-BINDING PROTEIN 1C/C-REPEAT-BINDING FACTOR 2; ERF = ETHYLENE RESPONSE FACTOR, PP2CA = PROTEIN PHOSPHATASE 2CA; PUMP4 = PLANT UNCOUPLING MITOCHONDRIAL PROTEIN 4; RAP2-13 (RAP2.4/WIND) = RELATED TO AP2 13; SZF1 = SALT-INDUCIBLE ZINC-FINGER; ZAT10 (STZ) = ZINC FINGER PROTEIN 10 (SALT TOLERANCE ZINC FINGER). Analysis was done using GeneMania (Warde-Farley *et al.*, 2010).

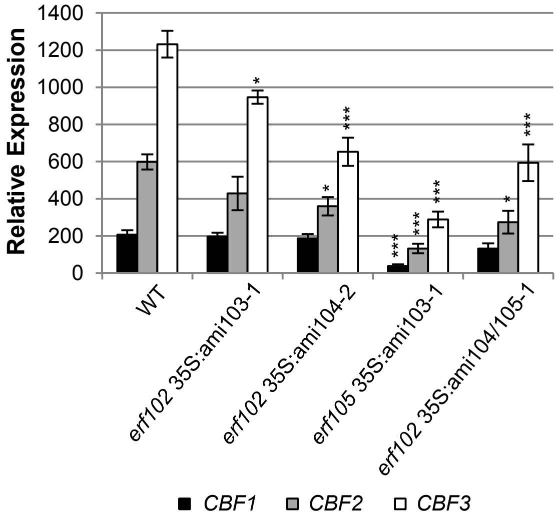


**Figure S6. Expression of selected cold-responsive genes in lines with reduced *ERF102* to *ERF105* expression.** Relative expression of *CBF1*, *CBF2* and *CFB3* genes in lines with reduced *ERF102* to *ERF105* after 4 hours of cold treatment at 4 °C. Transcript levels of wild-type samples under control conditions were set to 1 ($n \geq 4$). Asterisks indicate significant differences to the wild type (*, $p < 0.05$; ***, $p < 0.001$). Error bars represent SE.

**Table S1. Sequences of primers used for cloning.** Small letters in the primer sequences indicate the integrated *attB4*- or *attB1*-sites for cloning DNA fragments into the vector pDONR P4-P1R. Small italic letters in the primer sequences indicate the integrated *attB1*- or *attB2*-sites for cloning DNA fragments into the vector pDONR221. Underlined letters are the nucleotides added to keep the sequence in the right frame.

| **Amplification of** | **Primer sequences (5'–3')** |
|---|---|
| Promoter | F: ggggacaactttgtatagaaaagttgCGTTGATTCATTCTACAAACCAG |
| of *ERF102* (AT5G47230) | R: ggggactgctttttgtacaaacttgTGATAAAATTTTCAAAAAGC |
| Promoter | F: ggggacaactttgtatagaaaagttgGTTGTGGATTCTGGCATTTG |
| of *ERF103* (AT4G17490) | R: ggggactgctttttgtacaaacttgTTTGGAGGAAACAGAGAATTG |
| Promoter | F: ggggacaactttgtatagaaaagttgTGATGAGTGGTCGCTTCTTT |
| of *ERF104* (AT5G61600) | R: ggggactgctttttgtacaaacttgCTTCACTCTACTTGATTGAC |
| *ERF102* (AT5G47230) | F: *ggggacaagtttgtacaaaaaagcaggct*CAATGGCGACTCCTAACGAAG |
| for N-terminal fusion | R: *ggggaccactttgtacaagaaagctgggt*GTAATCAAACAACGGTCAACTGG |
| *ERF103* (AT4G17490) | F: *ggggacaagtttgtacaaaaaagcaggct*CGGAAATGGCTACACCAAACGAA |
| for N-terminal fusion | R: *ggggaccactttgtacaagaaagctgggt*CTCAAACAACGGTCAATTGTG |
| *ERF104* (AT5G61600) | F: *ggggacaagtttgtacaaaaaagcaggct*CGATGGCAACTAAACAAGAAGCTTTA |
| for N-terminal fusion | R: *ggggaccactttgtacaagaaagctgggt*CGTTTTAAGTGACGGAGATAACGGAAA |
